## Supplementary Figures for "Epimutations driven by RNAi or heterochromatin evoke transient antimicrobial drug resistance in fungi"

**Figure S1. Son et al.**

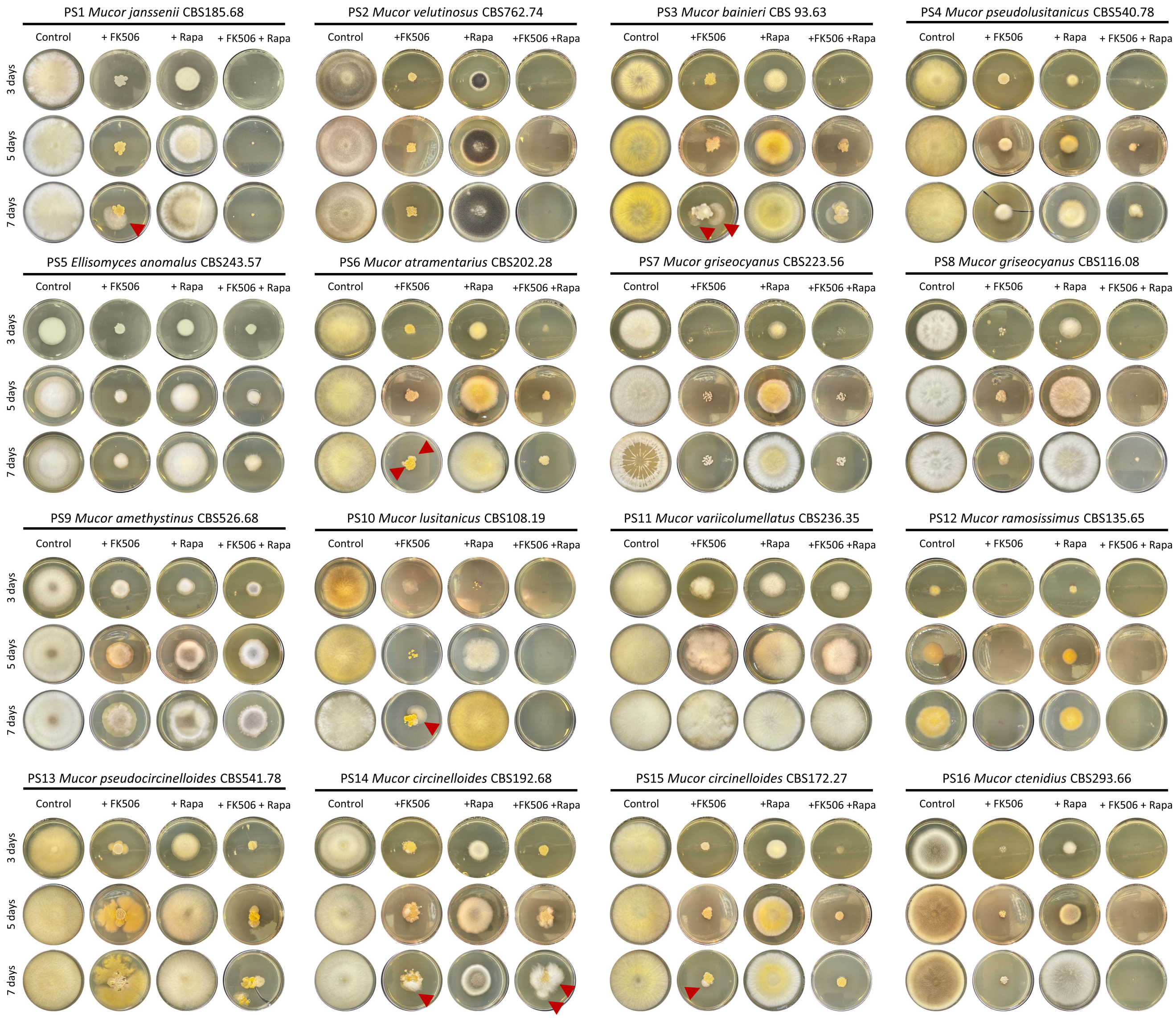

**Figure S2. Son et al.**

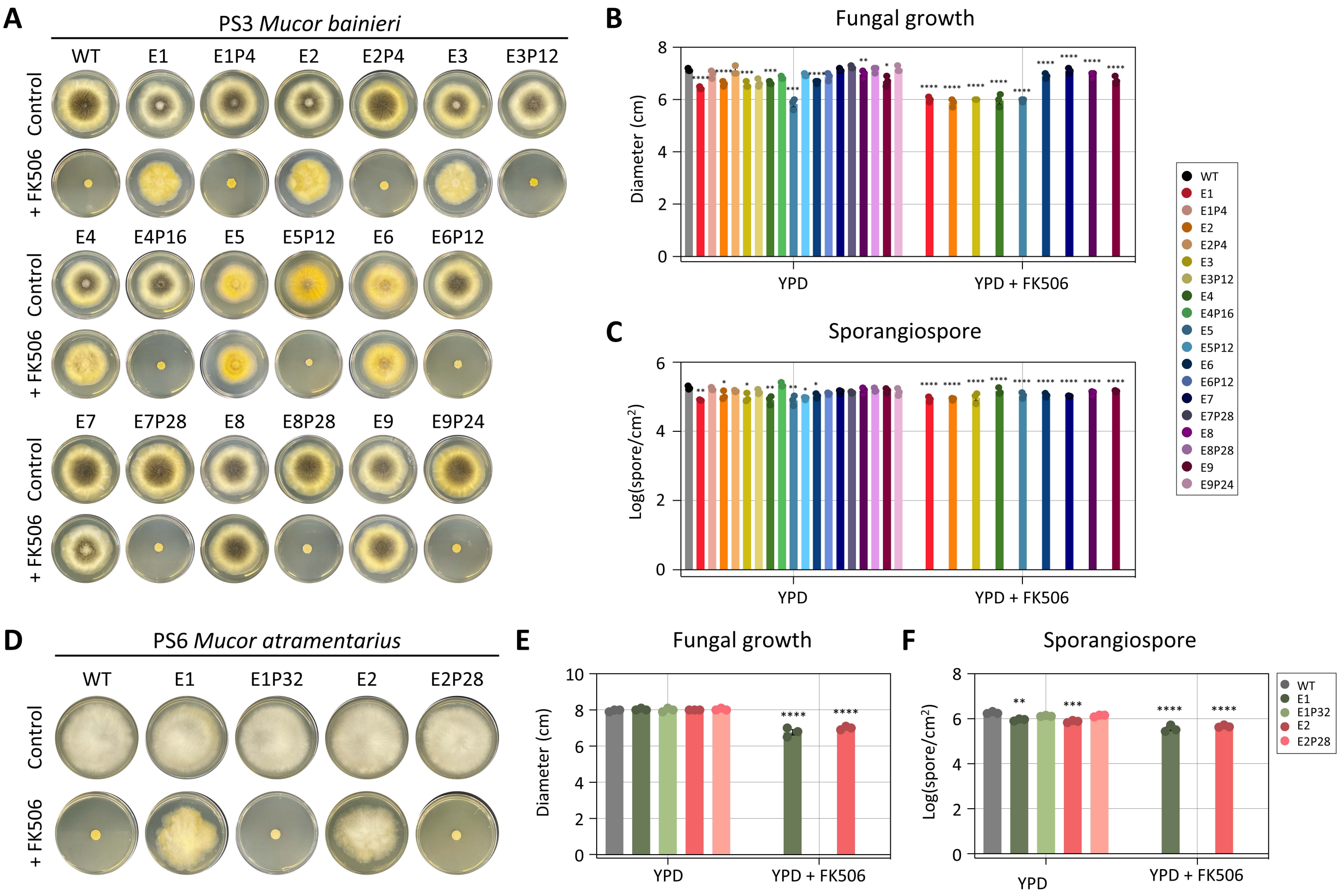

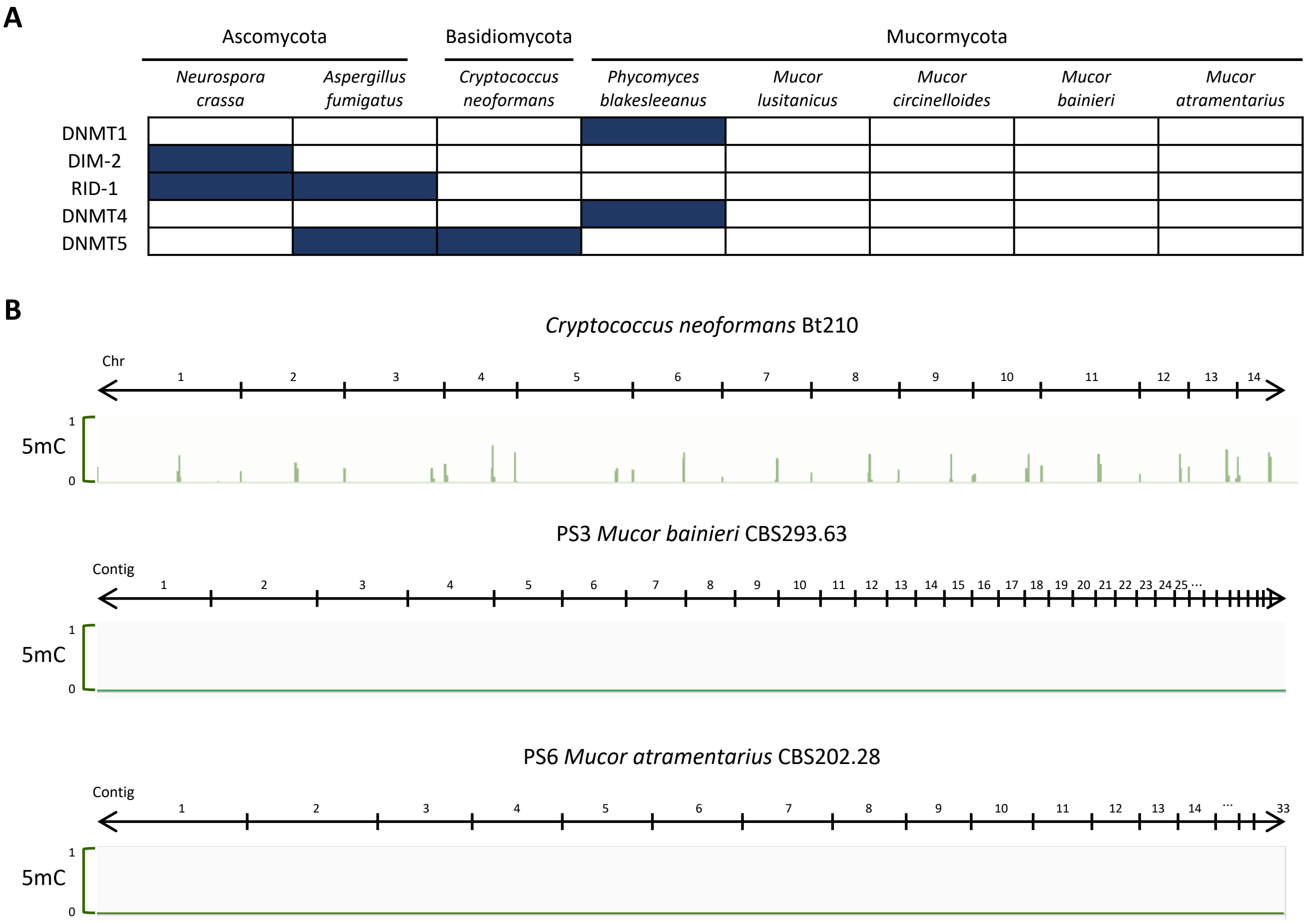
A
Ascomycota
Basidiomycota
Mucormycota
Neurospora crassa
Aspergillus fumigatus
Cryptococcus neoformans
Phycomyces blakesleeanus
Mucor lusitanicus
Mucor circinelloides
Mucor bainieri
Mucor atramentarius
DNMT1
DIM-2
RID-1
DNMT4
DNMT5
B
Cryptococcus neoformans Bt210
Chr
1 2 3 4 5 6 7 8 9 10 11 12 13 14
5mC
1
0
PS3 Mucor bainieri CBS293.63
Contig
1 2 3 4 5 6 7 8 9 10 11 12 13 14 15 16 17 18 19 20 21 22 23 24 25 …
5mC
1
0
PS6 Mucor atramentarius CBS202.28
Contig
1 2 3 4 5 6 7 8 9 10 11 12 13 14 … 33
5mC
1
0

**Figure S4. Son et al.**

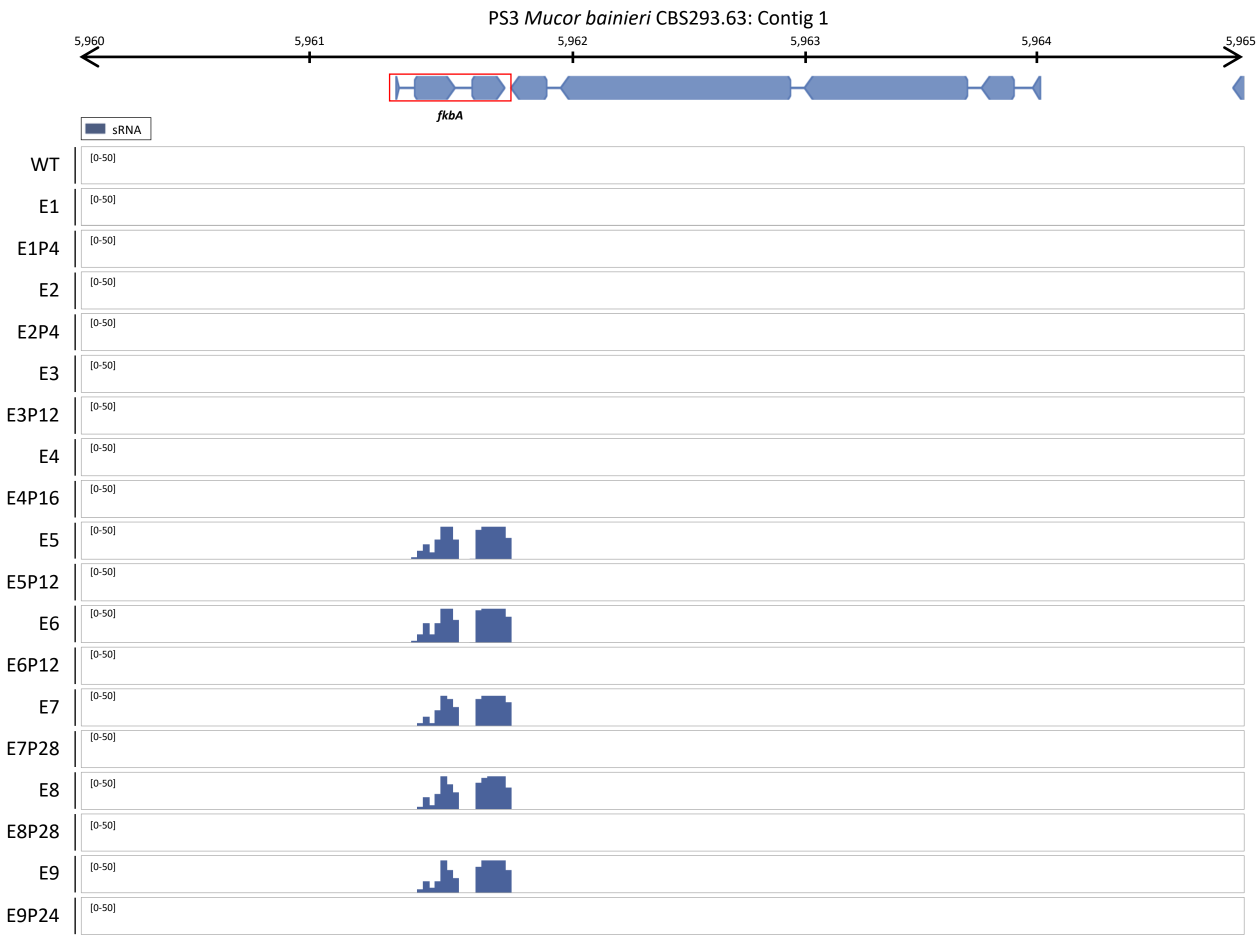

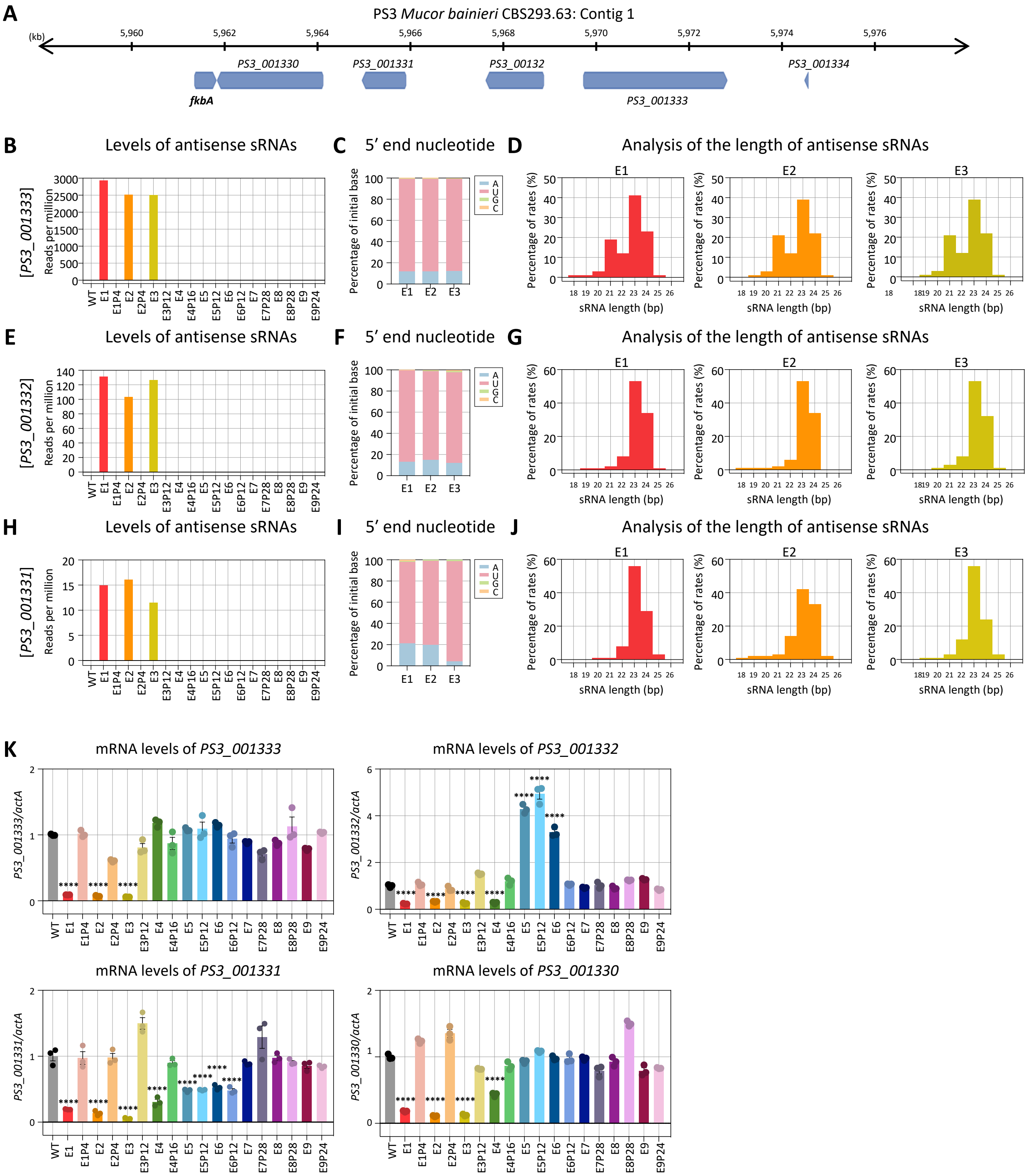
A
PS3 Mucor bainieri CBS293.63: Contig 1
(kb)
5,960
5,962
5,964
5,966
5,968
5,970
5,972
5,974
5,976
PS3_001330
PS3_001331
PS3_00132
PS3_001334
fkbA
PS3_001333
B
Levels of antisense sRNAs
[PS3_001333]
Reads per million
C
5' end nucleotide
Percentage of initial base
D
Analysis of the length of antisense sRNAs
E1
E2
E3
Percentage of rates (%)
sRNA length (bp)
E
Levels of antisense sRNAs
[PS3_001332]
F
5' end nucleotide
G
Analysis of the length of antisense sRNAs
H
Levels of antisense sRNAs
[PS3_001331]
I
5' end nucleotide
J
Analysis of the length of antisense sRNAs
K
mRNA levels of PS3_001333
PS3_001333/actA
mRNA levels of PS3_001332
PS3_001332/actA
mRNA levels of PS3_001331
PS3_001331/actA
mRNA levels of PS3_001330
PS3_001330/actA
WT
E1
E1P4
E2
E2P4
E3
E3P12
E4
E4P16
E5
E5P12
E6
E6P12
E7
E7P28
E8
E8P28
E9
E9P24

### Figure S6. Son et al.

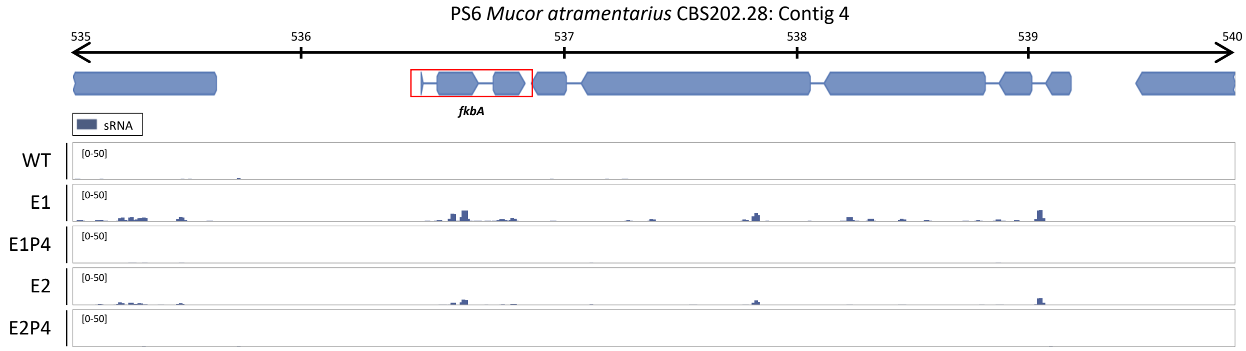

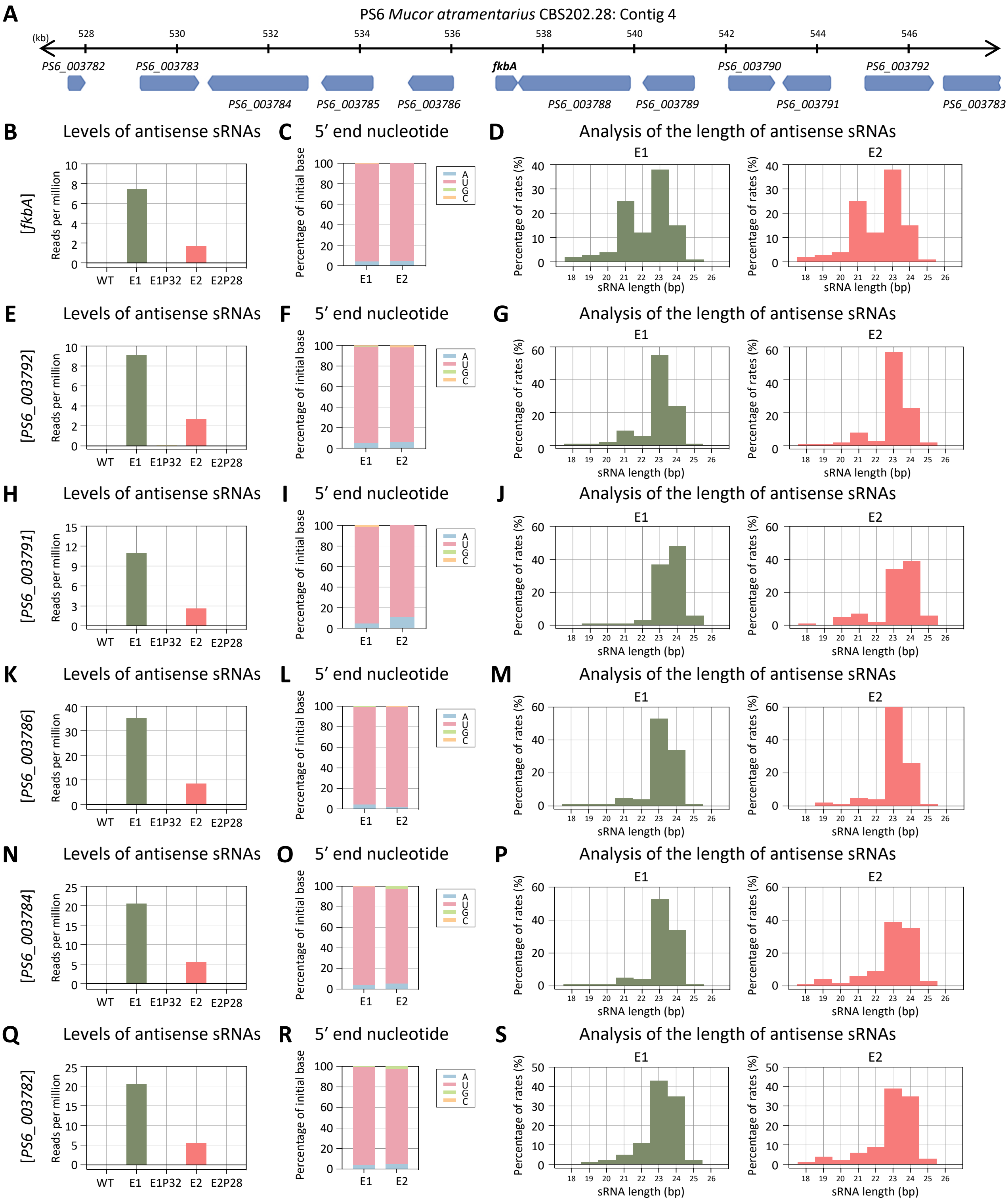
A
PS6 *Mucor atramentarius* CBS202.28: Contig 4
(kb)
528
530
532
534
536
538
540
542
544
546
PS6_003782
PS6_003783
PS6_003784
PS6_003785
PS6_003786
fkbA
PS6_003788
PS6_003789
PS6_003790
PS6_003791
PS6_003792
PS6_003783
B Levels of antisense sRNAs
C 5' end nucleotide
D Analysis of the length of antisense sRNAs
[fkbA]
E Levels of antisense sRNAs
F 5' end nucleotide
G Analysis of the length of antisense sRNAs
[PS6_003792]
H Levels of antisense sRNAs
I 5' end nucleotide
J Analysis of the length of antisense sRNAs
[PS6_003791]
K Levels of antisense sRNAs
L 5' end nucleotide
M Analysis of the length of antisense sRNAs
[PS6_003786]
N Levels of antisense sRNAs
O 5' end nucleotide
P Analysis of the length of antisense sRNAs
[PS6_003784]
Q Levels of antisense sRNAs
R 5' end nucleotide
S Analysis of the length of antisense sRNAs
[PS6_003782]
Reads per million
WT
E1
E1P32
E2
E2P28
Percentage of initial base
A
U
G
C
E1
E2
Percentage of rates (%)
sRNA length (bp)
18
19
20
21
22
23
24
25
26

**Figure S8. Son et al.**

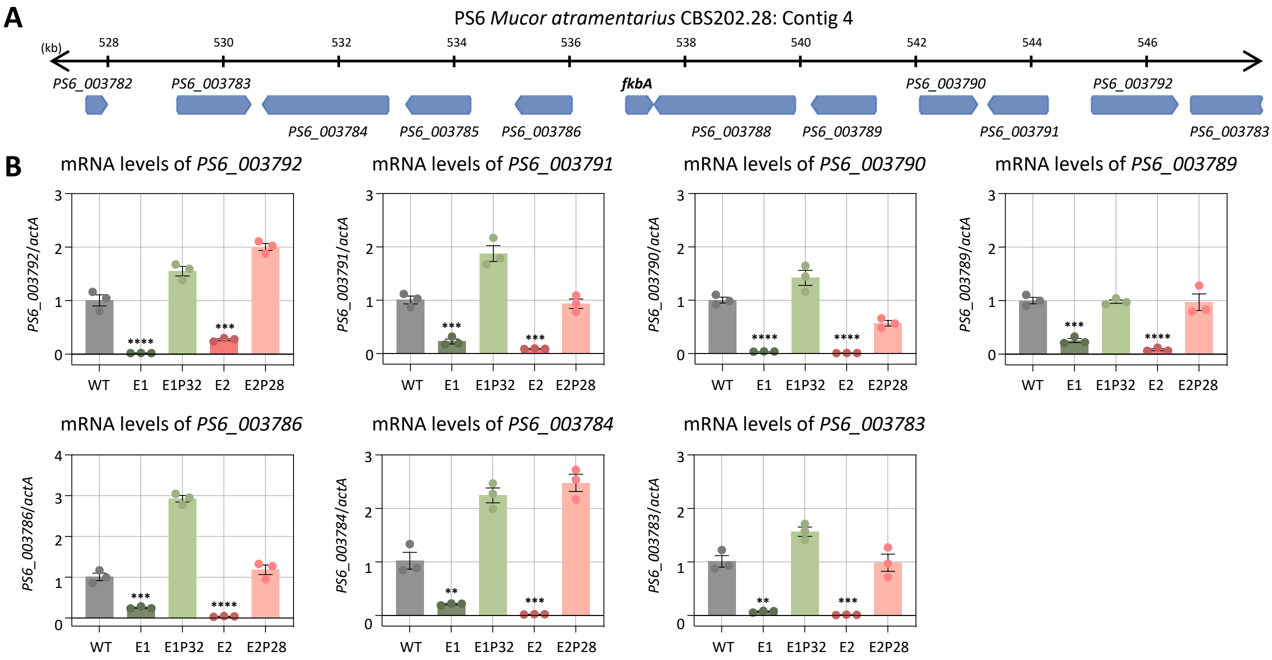

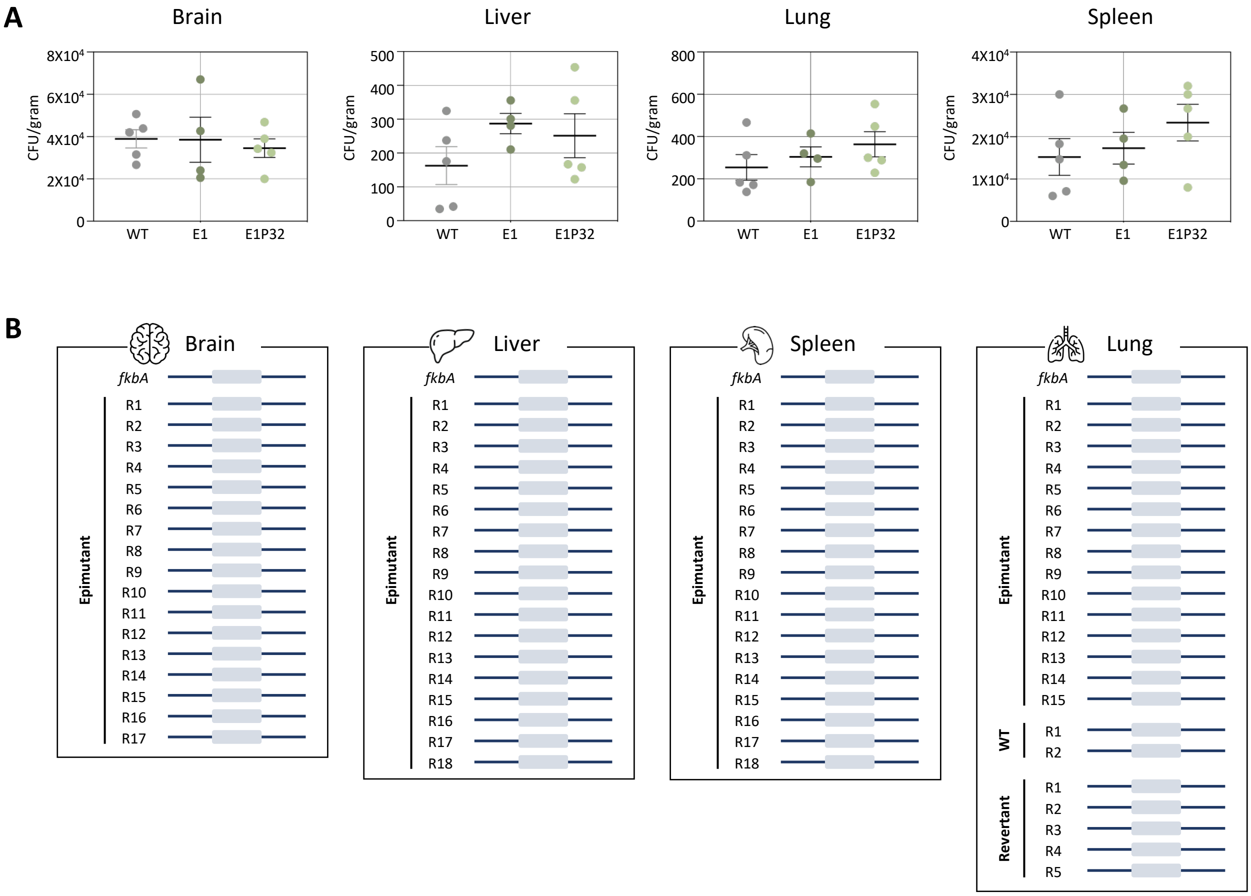
A
Brain
Liver
Lung
Spleen
CFU/gram
8X10^4
6X10^4
4X10^4
2X10^4
0
500
400
300
200
100
800
600
3X10^4
1X10^4
WT
E1
E1P32
B
Brain
Liver
Spleen
Lung
fkbA
Epimutant
R1
R2
R3
R4
R5
R6
R7
R8
R9
R10
R11
R12
R13
R14
R15
R16
R17
R18
WT
Revertant

**Figure S10. Son et al.**

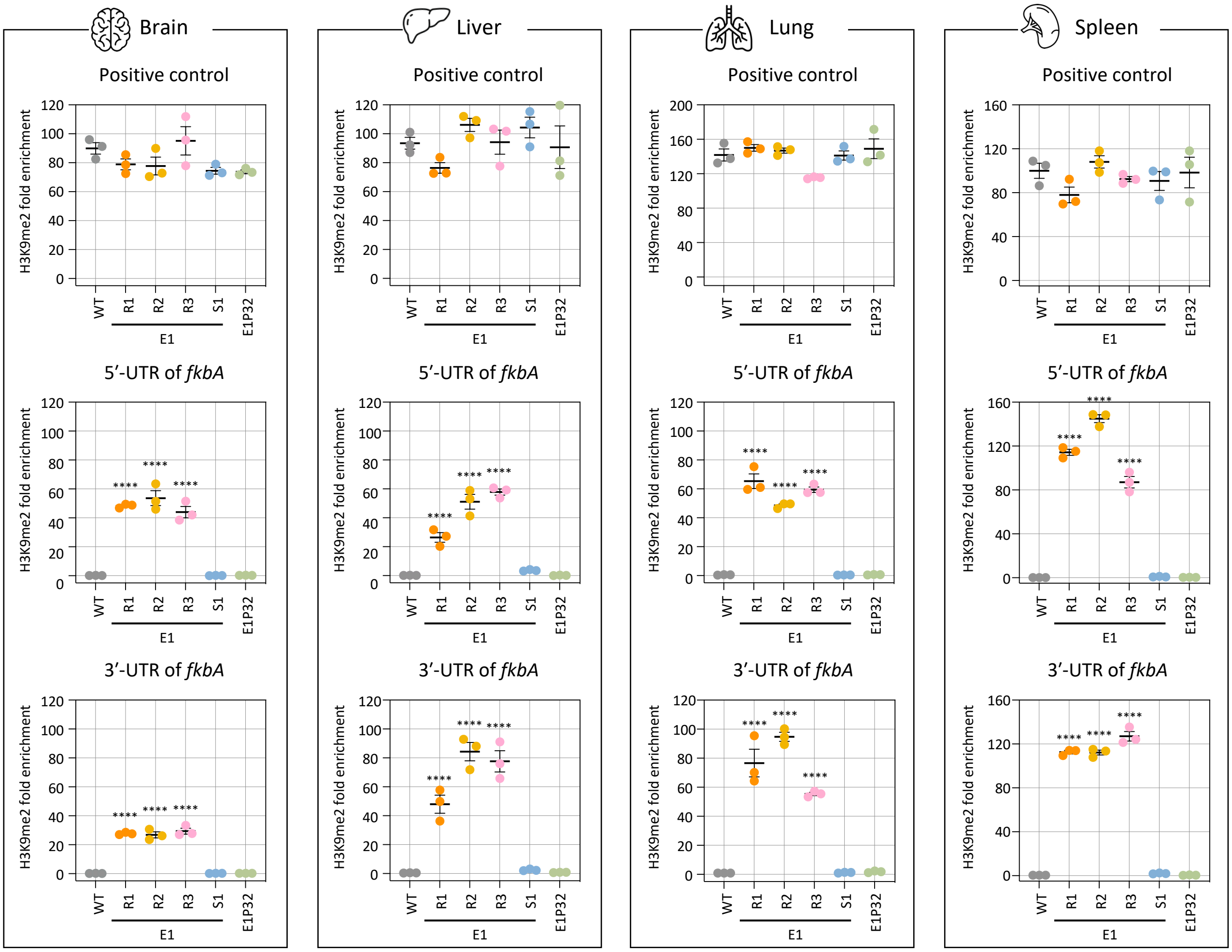

**Figure S11. Son et al.**

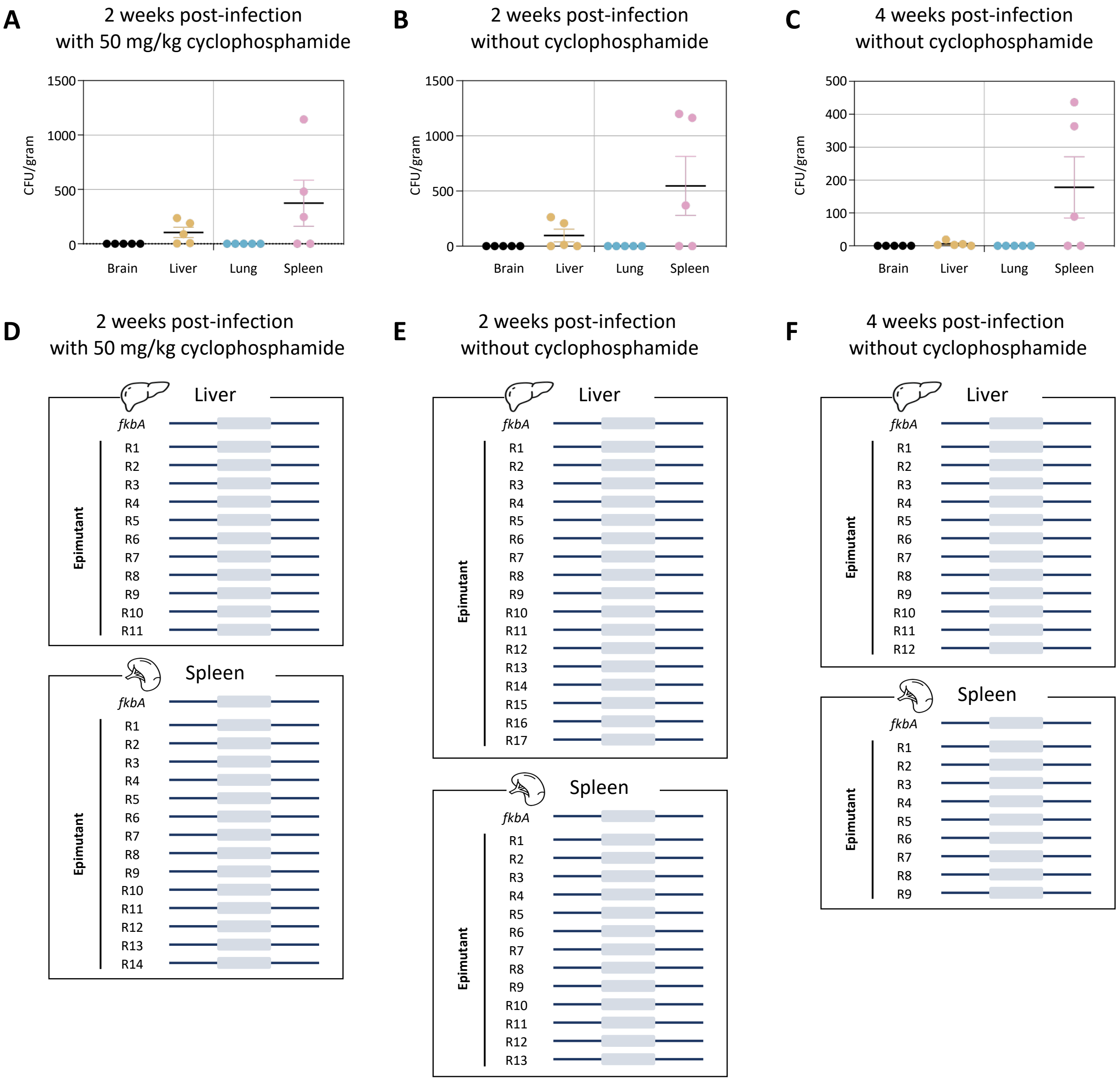

### Figure S12. Son et al.

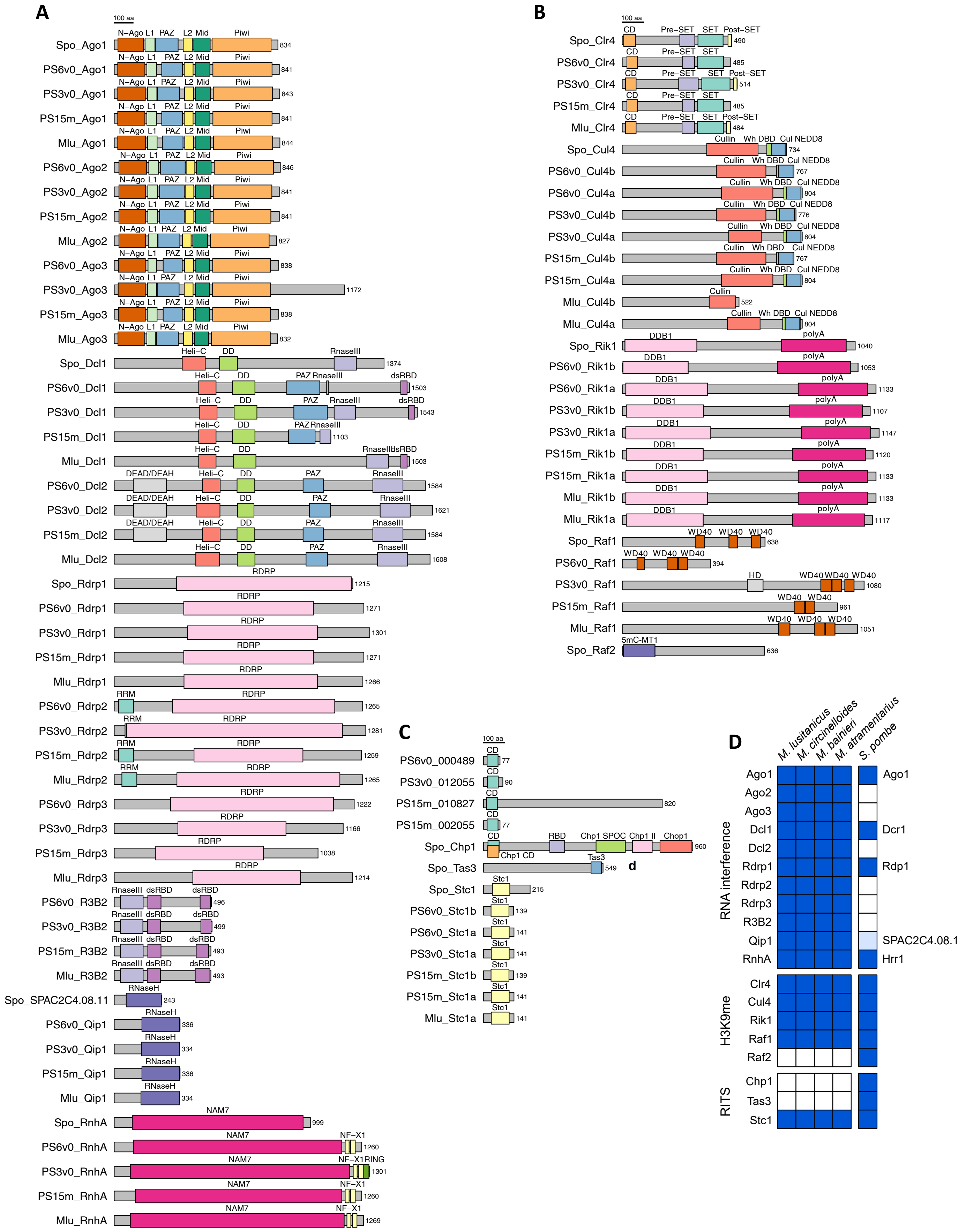

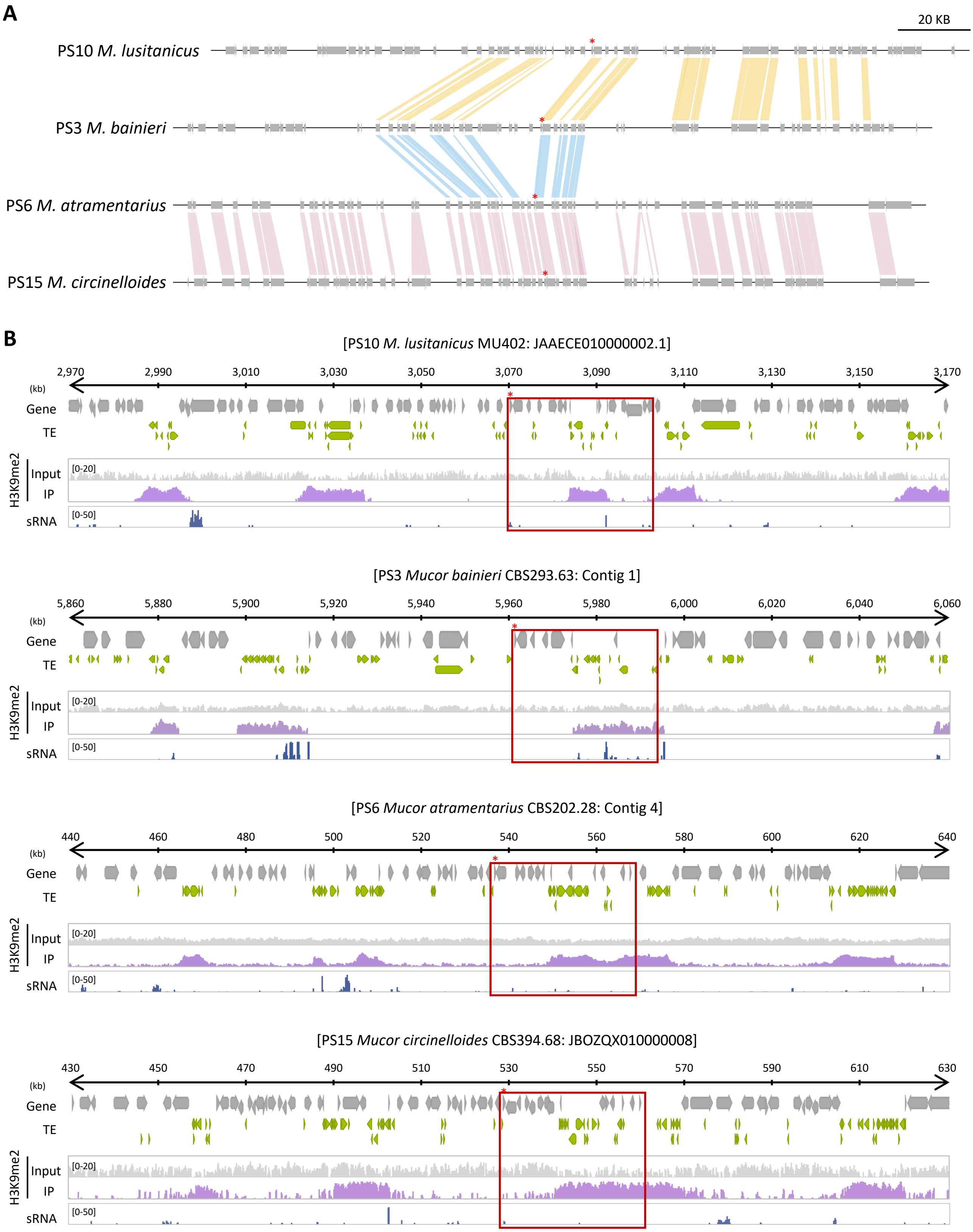
A
20 KB
PS10 M. lusitanicus
PS3 M. bainieri
PS6 M. atramentarius
PS15 M. circinelloides
B
[PS10 M. lusitanicus MU402: JAAECE010000002.1]
2,970
2,990
3,010
3,030
3,050
3,070
3,090
3,110
3,130
3,150
3,170
(kb)
Gene
TE
H3K9me2
Input
IP
[0-20]
sRNA
[0-50]
[PS3 Mucor bainieri CBS293.63: Contig 1]
5,860
5,880
5,900
5,920
5,940
5,960
5,980
6,000
6,020
6,040
6,060
[PS6 Mucor atramentarius CBS202.28: Contig 4]
440
460
480
500
520
540
560
580
600
620
640
[PS15 Mucor circinelloides CBS394.68: JBOZQX010000008]
430
450
470
490
510
530
550
570
590
610
630

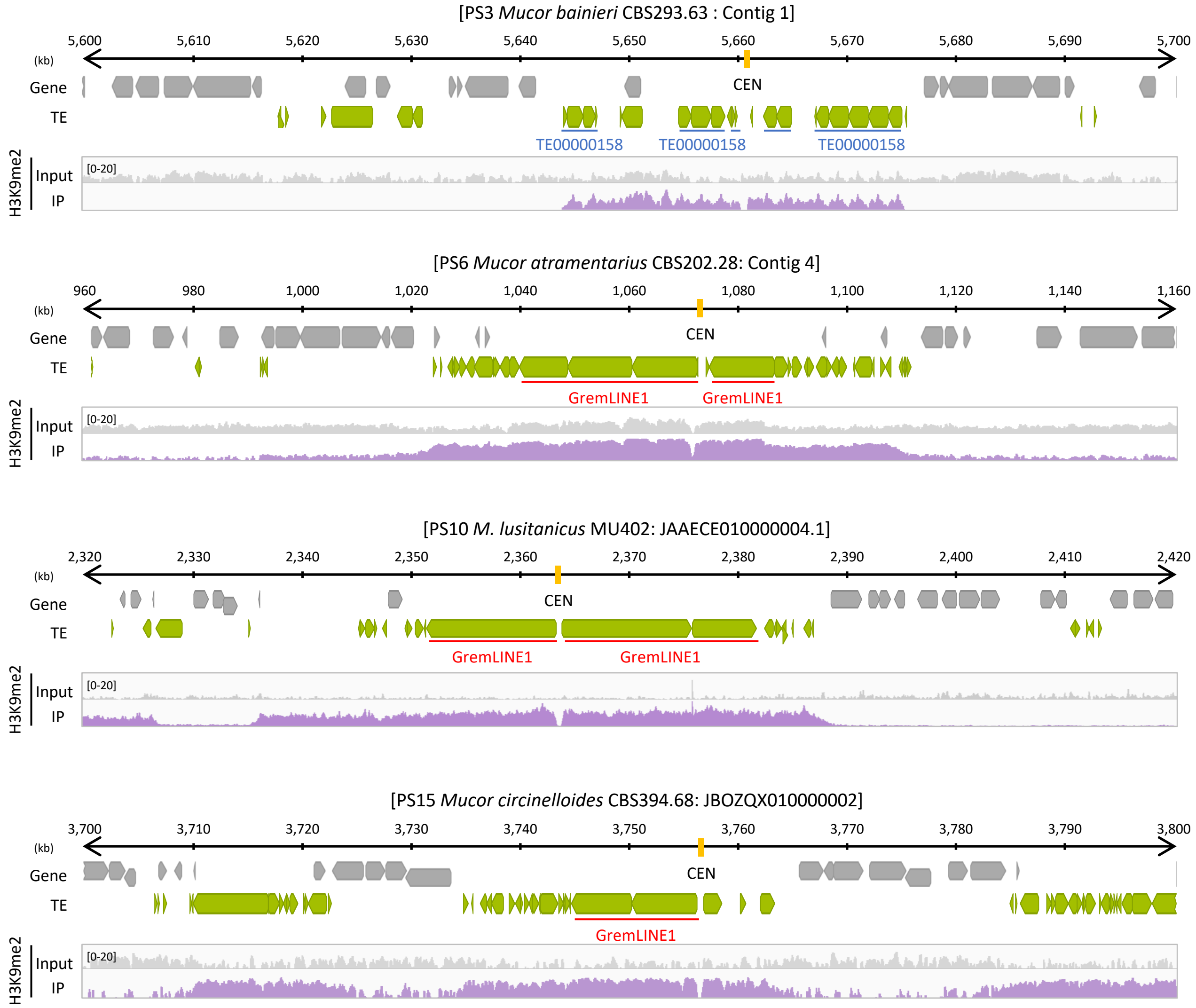
[PS3 *Mucor bainieri* CBS293.63 : Contig 1]
5,600
5,610
5,620
5,630
5,640
5,650
5,660
5,670
5,680
5,690
5,700
(kb)
Gene
TE
CEN
TE00000158
TE00000158
TE00000158
H3K9me2
Input
IP
[0-20]
[PS6 *Mucor atramentarius* CBS202.28: Contig 4]
960
980
1,000
1,020
1,040
1,060
1,080
1,100
1,120
1,140
1,160
(kb)
Gene
TE
CEN
GremLINE1
GremLINE1
H3K9me2
Input
IP
[0-20]
[PS10 *M. lusitanicus* MU402: JAAECE010000004.1]
2,320
2,330
2,340
2,350
2,360
2,370
2,380
2,390
2,400
2,410
2,420
(kb)
Gene
TE
CEN
GremLINE1
GremLINE1
H3K9me2
Input
IP
[0-20]
[PS15 *Mucor circinelloides* CBS394.68: JBOZQX010000002]
3,700
3,710
3,720
3,730
3,740
3,750
3,760
3,770
3,780
3,790
3,800
(kb)
Gene
TE
CEN
GremLINE1
H3K9me2
Input
IP
[0-20]
